# The Biosynthetic Costs of Amino Acids at the Base of the Food Chain Determine Their Use in Higher-order Consumer Genomes

**DOI:** 10.1101/2021.11.03.467059

**Authors:** Javier Gómez Ortega, Sonika Tyagi, Christen K. Mirth, Matthew D. W. Piper

## Abstract

Dietary nutrient composition is essential for shaping important fitness traits and behaviours. Many organisms are protein limited and for *Drosophila melanogaster*, this limitation manifests at the level of the single most limiting essential Amino Acid (AA) in the diet. The identity of this AA and its effects on female fecundity is readily predictable by a procedure called exome matching in which the sum of AAs encoded by a consumer’s exome is used to predict the relative proportion of AAs required in its diet. However, the exome matching calculation does not weight AA contributions to the overall profile by protein size or expression. Here we update the exome matching calculation to include these weightings. Surprisingly, although nearly half of the transcriptome is differentially expressed when comparing male and female flies, we found that creating transcriptome-weighted exome matched diets for each sex did not enhance their fecundity over that supported by exome matching alone. These data indicate that while organisms may require different amounts of dietary protein across conditions, the relative proportion of the constituent AAs remains constant. Interestingly, we also found remarkable conservation of exome matched AA profiles across taxa and that the composition of these profiles could be explained by the metabolic costs of microbial AA synthesis. Thus, it appears that bioenergetic constraints amongst autotrophs shape the relative proportion of AAs that are available across trophic levels and that that this constrains biomass composition.

## INTRODUCTION

Nutrition is one of the most important environmental determinants of evolutionary fitness; it supplies organisms with energy and the building blocks they require for growth, reproduction, and somatic maintenance (Simpson and Raubenheimer, 2012). However, the natural availability of food and its nutritional qualities vary and inevitably differ from the consumer’s needs (Denno and Fagan, 2003; Simpson and Raubenheimer, 2012; Wilder *et al*., 2013). As such, evolutionary fitness is constrained by the divergence between nutrient demand and their availability. Because of this, optimising nutrition to enhance growth, reproduction, and health is of major interest from both a fundamental biology and a commercial perspective.

Among the main components in the diet, protein is the limiting nutrient for the growth and reproduction of many organisms. It is, therefore, a principal constraint on evolutionary fitness (Simpson and Raubenheimer, 2012; Solon-Biet *et al*., 2015; Simpson *et al*., 2017; Jang and Lee, 2018). For example, the abundance of protein-rich food has been shown to increase population size or stimulate body growth of birds such as the galah (*Eolophus roseicapilus*) or the goldfinch (*Carduelis carduelis*) and mammals like the house mouse (*Mus musculus*) or several species of squirrels (*Sciurus and Tamiasciurus spp*.) (Smith, 1968, 1991; Nixon, 1970; Nixon, McClain and Donohoe, 1975; Glück, 1988; Keith and Bell, 1989; White, 1993). In the fruit fly *Drosophila melanogaster*, we have found that female reproduction is reduced by decreasing overall dietary protein concentration (Bass *et al*., 2007; Piper *et al*., 2014, 2017). We also found that this protein limitation is determined by the concentration of the single most limiting essential Amino Acid (AA) in the diet, which can be identified by comparing the proportion of AAs that is available in food against the proportion of AAs encoded by the fly’s exome – a procedure we called exome matching (Bass *et al*., 2007; Piper *et al*., 2014, 2017). Evidence from our work, and that of others, indicates that exome matching may have broader application as protein limitation also occurs at the level of single AAs in other species (Lochmiller *et al*., 1995; Ramsay and Houston, 1998; Webb *et al*., 2005; Piper *et al*., 2017).

To predict limiting AAs, our exome matching protocol involves two steps. First, we calculate each AA’s relative abundance in every protein of an organism’s *in silico* translated exome. Second, we find the average proportional representation for each AA across all proteins encoded by the exome. This genome-wide averaged AA proportion can then be compared to the AA proportion in the food to identify the essential AA that is most underrepresented in the diet and thus predicted to be limiting. We demonstrated that supplementing the diets of flies and mice with the limiting AA that was identified in this way can improve growth and reproduction and modify feeding behaviour (Piper *et al*., 2017; Solon-Biet *et al*., 2019; Sjøberg *et al*., 2020). Thus, for every organism whose genome has been sequenced, exome matching can theoretically be used as a tool to guide precision nutrition for better health.

Although we showed exome matching to be biologically effective, its current implementation does not incorporate weightings for the substantial differences we know to exist in genes’ sizes and their degree of expression (Oshlack and Wakefield, 2009). Many studies have documented considerable differences in gene expression profiles when comparing transcriptomes between sexes, across life-history stages, or in response to biotic and abiotic stimuli (Hegedus *et al*., 2009; May and Zwaan, 2017; Camus, Piper and Reuter, 2019; Li *et al*., 2019; Moskalev *et al*., 2019). For instance, in *Drosophila*, more than 8,000 genes, representing at least 50% of the genome, have been reported to be differentially expressed when comparing adult males with fertilised females – an observation that is unsurprising given the much heavier anabolic burden of reproduction for females than for males (Camus, Piper and Reuter, 2019). Thus, we predicted that we could improve the precision of exome matching by incorporating weightings for gene expression changes and in doing so, would establish a new way of tailoring diets to match an organism’s individual AA demands for life-stage and health status. Here, we set out to test this prediction and, in doing so, uncover that there is a surprisingly small variation in the way genomes from evolutionarily distant organisms encode AA usage. These data indicate a fundamental energy constraint on body composition across taxa.

## RESULTS

### CALCULATING TRANSCRIPTOME-WEIGHTED, EXOME-MATCHED DIETARY AA PROPORTIONS

Our previous research demonstrated that female flies fed food containing exome-matched AA proportions (FLYAA) laid more eggs than flies on food with equivalent amounts of protein comprised of mismatched AA proportions (Piper *et al*., 2017). Although FLYAA demonstrably improved egg-laying, we hypothesised it could be further improved by weighting each gene’s contribution to the overall average by its length and expression level. We reasoned that although there is not a 1:1 association between transcription and translation, the transcriptome would be a good approximation for the expression weightings for two reasons. First, transcriptomics readily yields a more complete set of gene expression values than proteomics (Pible and Armengaud, 2015; Manzoni *et al*., 2018). And, second, if the availability of dietary AAs constrains organismal protein expression, whole genome proteomics would simply reflect the constraints of diet quality. In contrast gene expression values may indicate protein expression levels that could be achieved if dietary AA availability was not a constraint - i.e. better matched to requirements.

We downloaded transcriptome profiles of whole male and whole female flies from FlyAtlas 2 and modENCODE, and from these profiles we averaged the levels of gene expression for each sex (Robinson *et al*., 2013; Brown and Celniker, 2015). To make our new sex-specific, transcriptome-matched profiles, we first counted the number of each AA that is encoded by each protein isoform in the fly genome. We then weighted these AA counts by the average isoform relative transcript abundance (FPKM value; see Materials and Methods) found for male or female flies. For each AA, we then summed the weighted AA counts across all genes and used these values to compute each AA’s proportional representation across all expressed genes. These newly designed AA ratios for the sexes were labelled MALEAA and FEMALEAA, and these became the basis for new dietary AA profiles (Figure 1).

**Figure 1.**
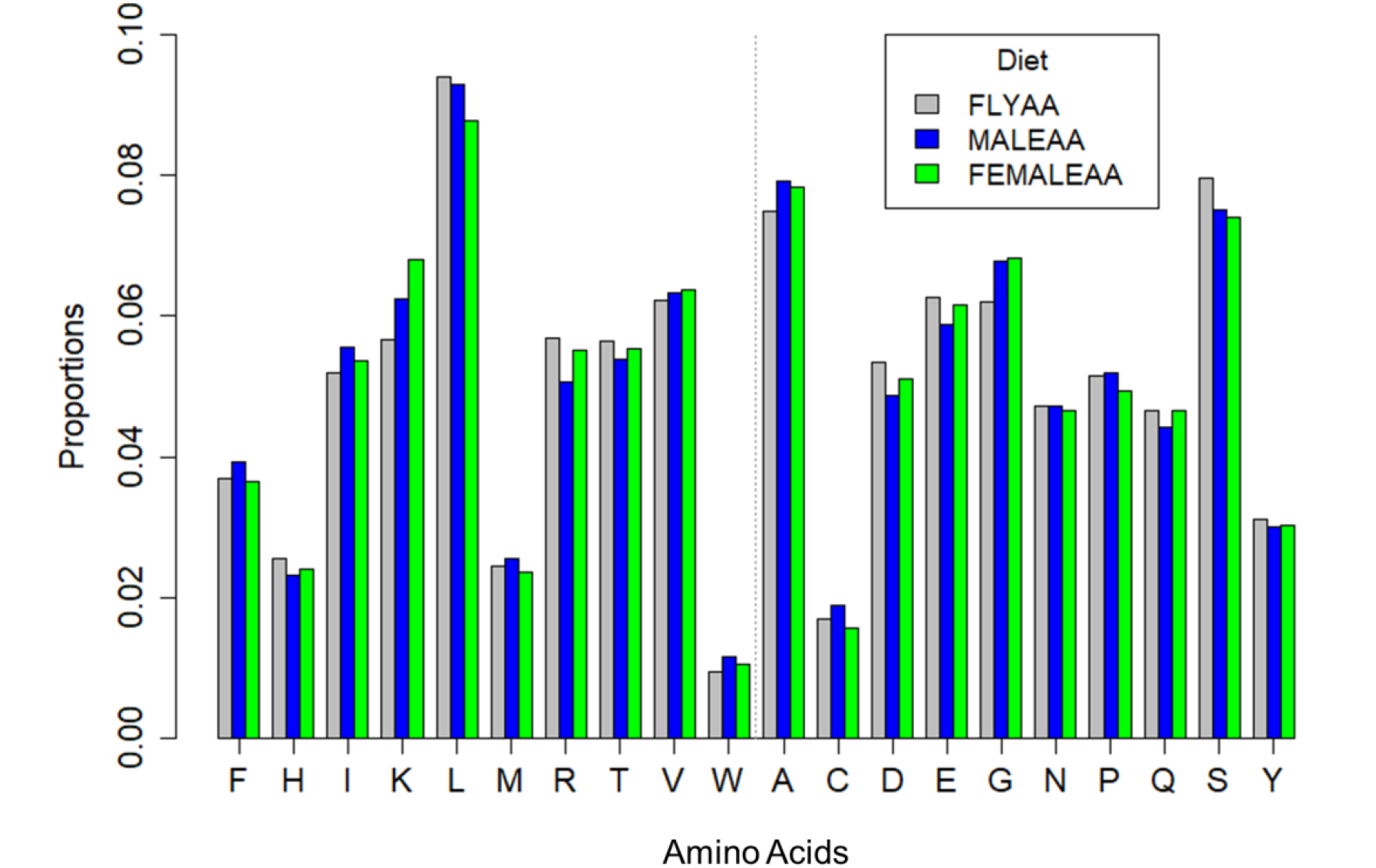
Comparison of the exome matched and transcriptome weighted AA ratios. Proportion of each AA in the exome matched (Piper *et al*., 2017) and transcriptome-weighted exome matched diets (MALEAA and FEMALEAA). In these diets, the average proportions of AAs have been generated after weighting each protein’s contribution by size and its average gene expression in male and female transcriptomes, respectively. The dotted grey line separates the essential (left) from the non-essential (right) AAs. IUPAC single-letter AA codes are shown.

### TRANSCRIPTOME-WEIGHTED EXOME MATCHING THE DIETARY AA PROFILE DOES NOT IMPROVE MALE FERTILITY OVER THAT ON AN EXOME MATCHED DIET

To test the effects of dietary AAs on male fertility, we used an assay in which males were challenged to inseminate females at maximum capacity, as this should deplete the males of sperm and/or seminal fluid and thus require them to be synthesising more from the dietary AAs they have available. To do this, we supplied singly housed males with ten new virgin females per day for seven days and counted how many of these females subsequently produced viable offspring. We found that while on the first day, males could inseminate 8 to 10 of these virgins, during the course of the assay, the number of females that each male could inseminate dropped to at least half of the number found for day one (Figure 2A) indicating that the males were indeed operating at maximum capacity in this assay.

**Figure 2.**
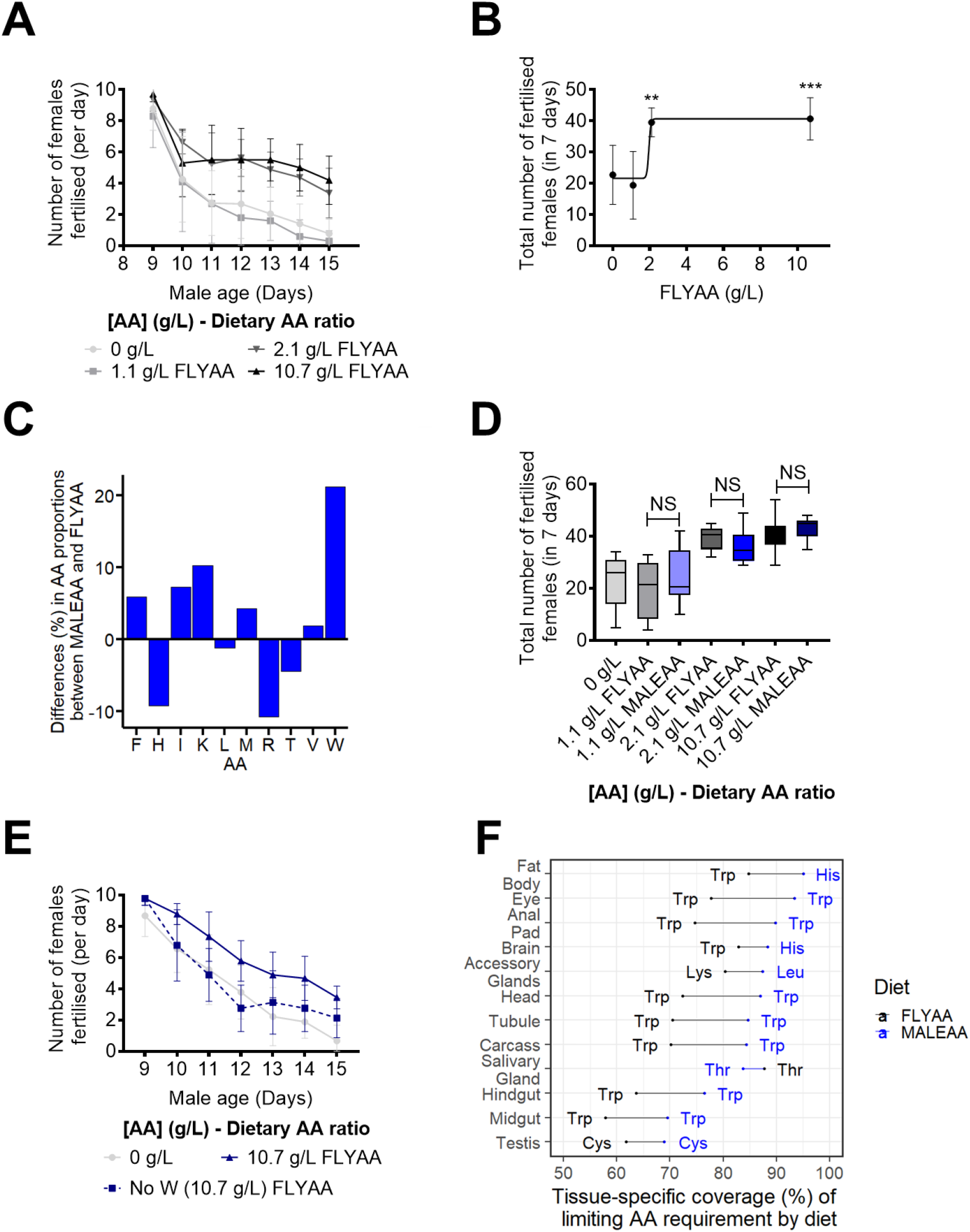
Male fecundity is modified by dietary AA concentration but not ratio. (A) The number of virgin female flies successfully fertilised by individual males during our seven-day assay. Male fecundity was reduced by decreasing dietary AA concentrations. AAs were provided in the exome matched proportion, FLYAA). Error bars represent the standard deviation. (B) The change in the cumulative number of females fertilised by males in response to dietary AA change could be modelled by a sigmoidal dose-response curve (R^2^ = 0.537; least-squares fit). (*** = P<0.001, in comparison with 0 g/L). Error bars represent the standard deviation. (C) Predicted difference in each essential AA when comparing the male transcriptome matched proportions (MALEAA), and the exome matched proportions (FLYAA). A positive difference indicates that the AA is more abundant in MALEAA than in the FLYAA. MALEAA should cover any essential AA deficiency of FLYAA, and thus, the relative increase in the concentration of the most limiting essential AA (Tryptophan, W, 20%) equals the potential increase in fecundity that could be achieved for flies fed with MALEAA. (D) Males fed with a diet containing a transcriptome (MALEAA) matched AA proportion did not differ from those fed the exome matched (FLYAA) diets for any concentration of AAs tested. Error bars represent the standard deviation. (E) The effect of tryptophan dropout from the diet on the daily capacity of males to fertilise females during a seven-day period. The removal of tryptophan from the diet caused a fast decay in fertilisation that matched that caused by the removal of all AAs. AAs were provided in the male transcriptome matched proportion, MALEAA. Error bars represent the standard deviation. (F) Coverage of the predicted dietary AA requirements by MALEAA and FLYAA when compared to the transcriptome weighted exome match proportion from each tissue in male flies. For each tissue, the x-axis displays the degree to which the limiting AA demand is met by the diets MALEAA (blue) and FLYAA (black). The closer to 100%, the better the diet covers the predicted tissue demand for AAs. For all tissues, except the salivary gland, MALEAA is predicted to be a better match for requirements than FLYAA. The predicted limiting AA for each tissue on each diet is indicated by the three letter AA codes.

To test if male fecundity changed in response to altered dietary protein levels, we performed the above assay on males that were maintained on food in which the AA proportion was fixed (FLYAA), but the total concentration was diluted from 10.7g/l (positive control; the level in our standard “rich” diet) to 2.1 g/l, 1.1 g/l and 0 g/l. Male fertility was significantly lower than the positive control when the flies were maintained on 0 g/l and 1.1 g/l AAs (P<0.001) (Figure 2A). Surprisingly, when the protein concentration was 2.1 g/l, male fertility increased to the level of flies maintained on the positive control condition (Figure 2A, 2B). Thus, maximum male fertility in our assay responded to dietary AA levels and only relatively small amounts were required to support maximal fertility. The response of male fecundity to dietary AA concentration could be modelled by a sigmoidal dose-response curve with an inflexion point somewhere between 1.1 – 2.1 g/l AA (Figure 2B).

If MALEAA represents the ideal proportion of AAs for male fecundity, we predict that males fed FLYAA would be tryptophan (trp, W) limited, and that for a fixed sum of AAs changing the proportion to MALEAA would yield a 20% increase in AA availability for reproduction (Figure 2C). However, when we compared the fecundity response of male flies kept on MALEAA and FLYAA at each of the dietary AA concentrations, we saw that AA ratio did not alter the number of females that were successfully inseminated, even under conditions where male fertility was clearly AA limited (1.1 g/L; Figure 2D).

A possible reason why MALEAA did not improve fecundity is that males might contain sufficient stores of tryptophan in body proteins that they can retrieve and use to overcome the limitation we predicted. If this were the case, our prediction of a 20% improvement in fecundity would be an overestimate. To assess this, we made another diet in which only tryptophan was omitted from the diet altogether. We evaluated the effect of this diet and found that it caused a significant reduction in fecundity compared to the positive control diet (10.7g/l). It was also equally as detrimental for fecundity as a diet without AAs, both in terms of the rate at which fertility fell, and the total number of females successfully fertilised during the assay (Figure 2E). Thus, dietary tryptophan is required to sustain male fecundity in this assay, and its requirement does not appear to be lessened due to the recovery of tryptophan stored in body tissue.

Another reason why MALEAA may not have improved fecundity over FLYAA is that the actual set of proteins required for male fecundity are only a subset of those included in our calculation and that MALEAA is actually a worse AA balance for the organs responsible for making the proteins required for fecundity. To investigate this possibility, we calculated transcriptome-weighted AA profiles for each tissue type in male flies using RNAseq data from FlyAtlas (Robinson *et al*., 2013). We then compared these tissue profiles to both the unweighted (FLYAA) and transcriptome weighted (MALEAA) dietary AA profiles and predicted each tissue’s limiting AA and the degree to which it is limiting. The data are expressed as a relative match where 0 indicates the complete absence of an essential AA and 100 represents that all dietary AAs are perfectly matched to the tissue-specific profile (Figure 2F). The data show that MALEAA is predicted to be a better match than FLYAA for the expression of the genes in each tissue, except for those in the salivary glands in which FLYAA is predicted to be a better match than MALEAA. Thus, we still predict that MALEAA would be an improved AA profile over FLYAA for male reproduction if transcriptome weighting the exome provided a superior prediction of dietary requirements. However, our tissue-specific analysis does reveal that MALEAA is predicted to confer a smaller improvement over FLYAA if only the profile of the testis (62% – 68%; 9.6% increase) or accessory glands (80% – 88%; 10% increase) matter for our assay of male fecundity. It is possible that this small degree of enhancement in male fecundity was beyond the sensitivity of our assay to be detected.

### TRANSCRIPTOME-WEIGHTED EXOME MATCHING THE DIETARY AA PROFILE DOES NOT IMPROVE FEMALE FECUNDITY OVER THAT ON AN EXOME MATCHED DIET

Our previous data indicate that female egg-laying is a reliable indicator of dietary AA composition, and typically has lower variability and greater sensitivity than the male fecundity assay we performed (Piper *et al*., 2017). We thus tested if FEMALEAA had a higher nutritional value than FLYAA to sustain female fecundity. Two-day-old mated females were placed on chemically defined diets, and the number of eggs they laid over the course of eight days was counted. In line with previous results (Piper *et al*., 2017), female egg production responded in a linear manner to increasing AA concentrations until at least 10.7 g/L (Figure 3A). This is consistent with dietary AAs quantitatively limiting female egg-laying, which we have previously shown to be due to the most limiting essential AA (Piper *et al*., 2017).

**Figure 3.**
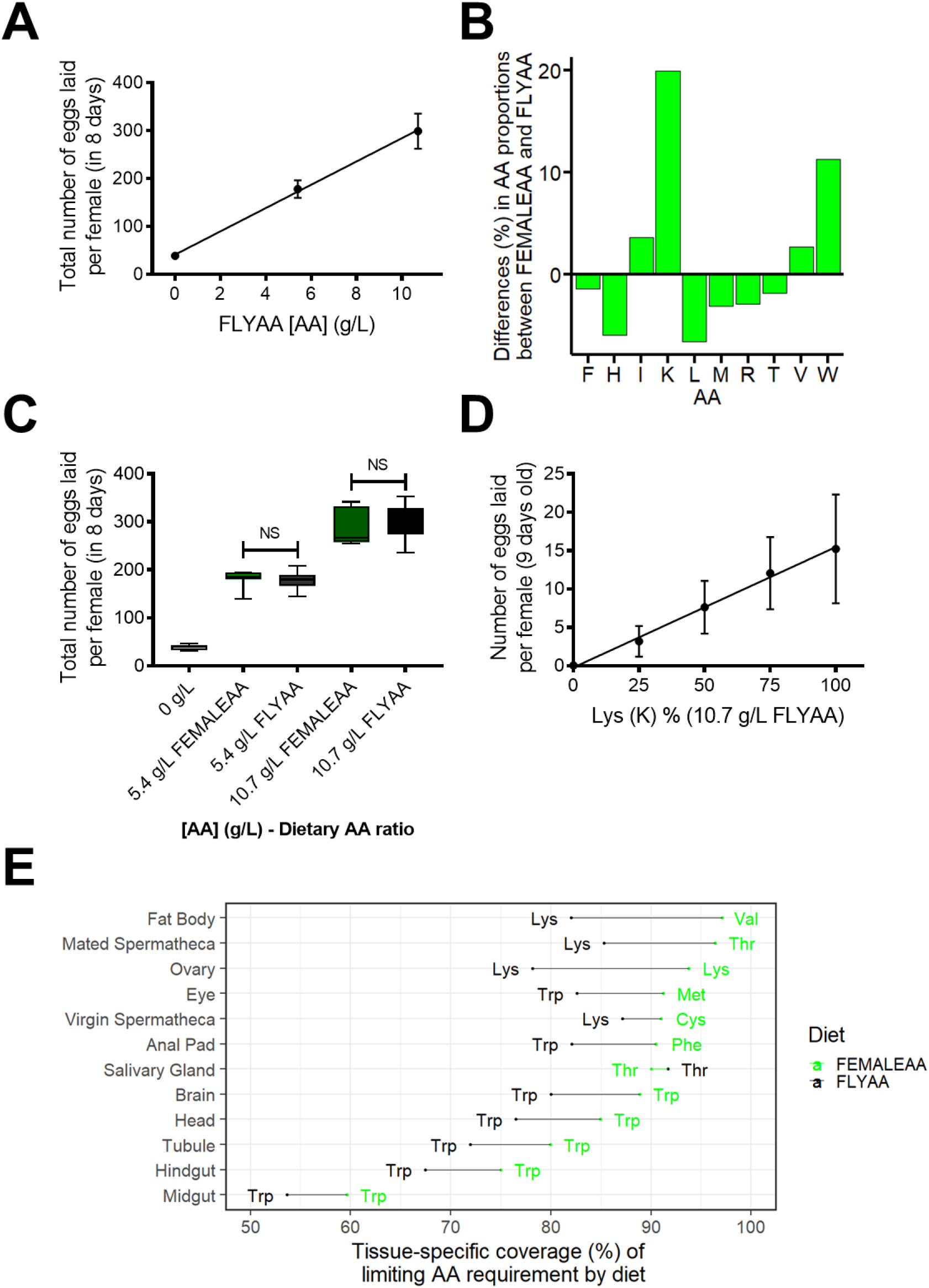
Female fecundity responses to changes in dietary AA ratio and concentration. (A) Total female fecundity showed a linear response to increasing dietary AA concentrations (FLYAA). R^2^ = 0.999 (B) Relative change in the molar concentration of each essential AA between the female transcriptome matched diet (FEMALEAA) and the exome matched diet (FLYAA). A positive difference indicates that the AA is more abundant in FEMALEAA than in FLYAA. The relative increase in the concentration of the most limiting essential AA (Lysine, K, 20%) equals the potential increase in fecundity that could be achieved for flies fed with FEMALEAA. (C) At each concentration of total AA, there was no difference in egg-laying output of females fed with transcriptome (FEMALEAA) or exome (FLYAA) matched diets. (D) Lysine dilution limits egg production in a linear manner. Percentages of Lysine concentration are relative to the standard Lysine concentration on FLYAA with a total AA concentration of 10.7 g/L. R2 = 0.995. (E) Coverage of the predicted dietary AA requirements of each tissue in female flies by FEMALEAA and FLYAA. Tissue dietary AA requirements are predicted from the tissue-specific transcriptomes. For each tissue, the x-axis displays the percent of the limiting AA demand covered by the diets FEMALEAA (green) and FLYAA (black). The closer to 100%, the better the diet meets the theoretical tissue AA demand. For all tissues, except the salivary gland, FEMALEAA is predicted to be a better match for requirements than FLYAA. The predicted limiting AA for each tissue on each diet is indicated by the three-letter AA codes.

We predicted that if the transcriptome-weighted diet (FEMALEAA) represented the actual AA requirements for females for egg-laying, lysine (K, Lys) would limit egg production for females feeding on the non-weighted AA ratio (FLYAA) (Figure 3B). By comparing the AA profile of FEMALEAA to that of FLYAA, we also predicted that egg production could be up to 20% higher when incorporating transcriptome weightings into the exome match profile (Figure 3B). However, FEMALEAA did not improve female fecundity output in comparison to FLYAA at either concentration of dietary AAs tested (Figure 3C). This included a concentration at which AAs clearly limited egg production (5.4 g/L) and so should be the most sensitive test of the change in availability of the most limiting AA.

We tested if the reason why female flies produced no more eggs on FEMALEAA than when on FLYAA is that they could supplement limiting dietary lysine by retrieving it from body protein reservoirs. If dietary lysine limits egg-laying, and the flies do not supplement it from body reserves, the flies should exhibit reduced egg production in proportion to the dilution of lysine in the diet. This is exactly what we observed when we reduced lysine only in an otherwise constant nutritional background containing 10.7 g/L FLYAA (Figure 3D). This demonstrates that lysine is both essential and limiting in FLYAA for female fecundity and indicates that lysine limitation is not lessened by flies retrieving it from stored body protein.

We also tested if FEMALEAA is not superior to FLYAA to support egg production because the flies use only a subset of the transcriptome to produce eggs. To do this, we assessed the match between each tissue-specific AA profile and FEMALEAA or FLYAA. Similar to the comparison we made for males, FEMALEAA was a better match than FLYAA to the transcriptome weighted AA proportions of every female tissue except for the salivary glands (Figure 3E). Furthermore, if we consider only the tissues most relevant to reproduction, the ovaries and fat body, FEMALEAA represents a better match, and to a similar extent as whole-body samples, to their transcriptome-weighted exome profiles than FLYAA (16% and 15% for ovaries and fat body respectively). These data support our prediction that FEMALEAA should improve dietary AA availability for egg production over that found in FLYAA.

### COMPARING GENOME-WIDE AA USAGE ACROSS TAXA INDICATES CONSTRAINTS FROM THE METABOLIC COSTS OF THEIR BIOSYNTHESIS

We have found that weighting the dietary exome matched AA ratio (FLYAA) by the average gene expression of male (MALEAA) or female (FEMALEAA) flies did not modify fly fecundity. This was surprising because the weightings incorporated changes in ∼50% of the expressed transcriptome. These data indicate that despite these large changes in gene expression, there are constraints on the degree to which genome-wide expression changes can modify the dietary AA requirements of our flies. One of the ways this could happen is if the male and female transcriptome-weighted AA profiles converge on a similar AA usage as the non-weighted (FLYAA) profile. To assess the degree of similarity of our three AA profiles (FEMALEAA, MALEAA and FLYAA), we compared them in the context of a null distribution of expression profiles. To generate this null distribution, we permuted the gene labels on the male and female transcriptome-wide expression data 20,000 times, thus generating a set of divergent but biologically realistic transcriptome-wide expression profiles. For each of these profiles, we then calculated the transcriptome-wide AA usage, and then calculated the Euclidian distance between each of the permuted AA profiles and the median permuted profile. This revealed that the AA proportions of FLYAA, FEMALEAA and MALEAA were more distant from the median transcriptome-weighted AA profile than most of the permutations (>99% for FEMALEAA, >92%, for MALEAA, and >97% for FLYAA in the context of FEMALEAA and >74% for FLYAA in the context of MALEAA), indicating that, compared with the null distribution, they all represented extreme examples of AA usage (Figure 4 A & B). Because Euclidian distance only measures the degree of divergence between profiles and not the direction of change (i.e. whether an AA is more or less abundant), we plotted the proportional representation of each AA for each AA profile (Figure 4C & D). When we did this, we found that for the AAs where MALEAA and FEMALEAA lay beyond the limits of the box plots, thus differing from the majority of permuted values, FLYAA tended to differ from the permuted values in the same way. Thus, the actual transcriptome weighted AA profiles that we found for males and females used AA in a manner more similar to the non-weighted AA profile (FLYAA) than the majority of the permuted profiles. This could indicate a constraint on AA usage that limits the degree to which the transcriptome can modify the consumer’s AA requirements.

**Figure 4.**
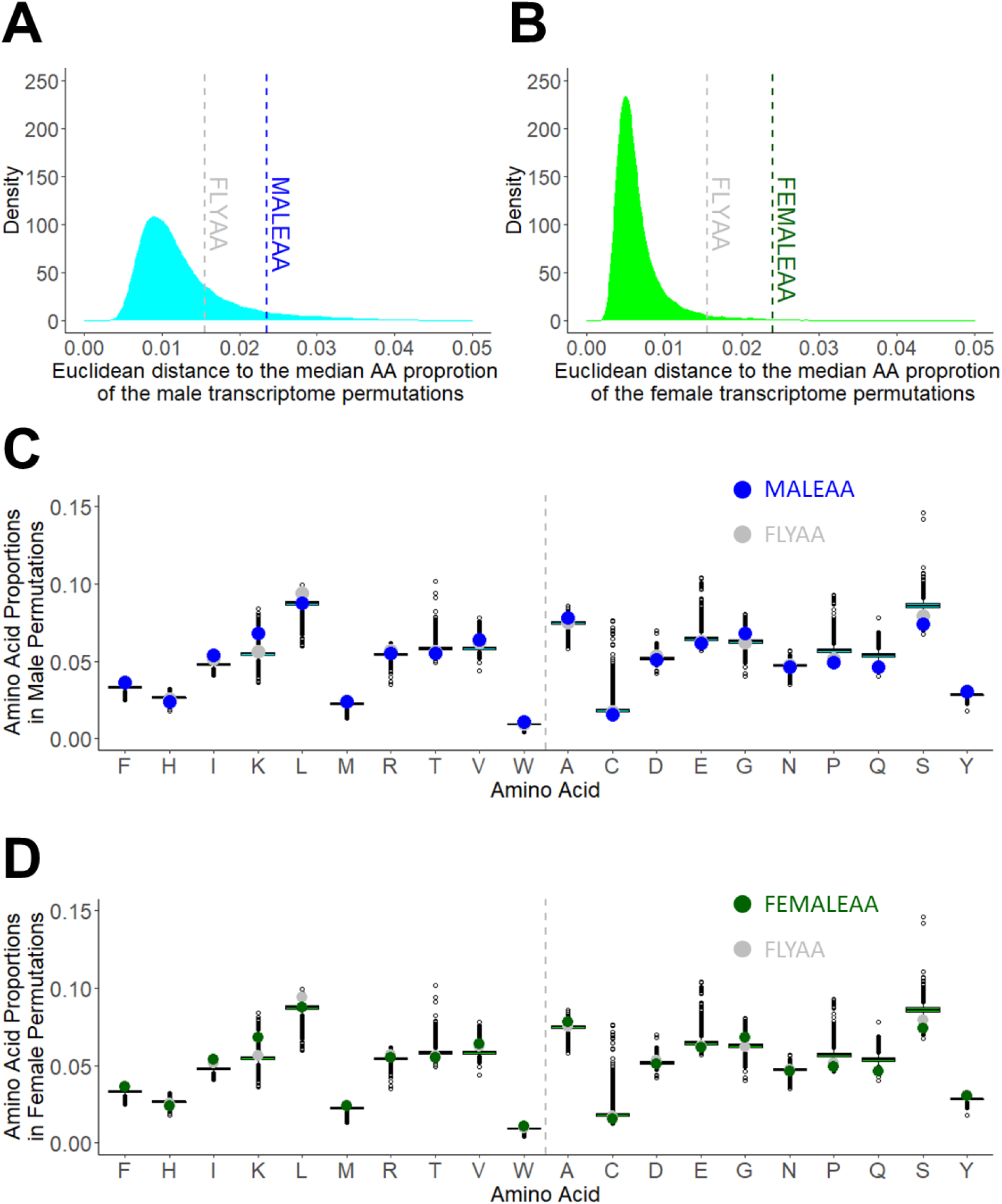
using permuted transcriptome-weighted AA profiles to compare the way in which the sex-specific transcriptome weighted profiles differ from exome matching. (A, B) We permuted the gene labels in our male (A) and female (B) transcriptome data 20,000 times and calculated a new transcriptome-weighted AA profile for each permutation. We then calculated the Euclidean distances from the median AA proportion of the permuted profiles to each permutation from the male (cyan) (A) and female (light green) (B) transcriptome data as well as the distance to MALEAA (dashed blue line) (A), FEMALEAA (dashed green line) (B) and FLYAA (dashed grey line) (A,B). (C, D) The relative proportion of each AA for all permuted profiles as well as the proportions found for MALEAA (blue) (C), FEMALEAA (green) (D), and FLYAA (grey) (C, D). The vertical grey dashed line separates essential (left) from non-essential (right) AAs. The median AA values are represented by a horizontal line dividing the boxes. The boxes represent the interquartile range (50% of the permuted values).

For organisms to maximise efficiency when using the limited amounts of dietary protein that are available, we anticipate that any constraints on AA usage in an individual organism may also be conserved across species. In particular, when moving up trophic levels, we expect that AA heterotrophs, which are higher-level consumers and cannot synthesise all AAs *de novo*, may contain very similar AA profiles to the diet they consume because they are confined to using AAs in the proportions found in their food. By contrast, AA autotrophs, which can produce all 20 protein-coding AAs, might have more divergent profiles. To assess this, we compared the exome-matched AA proportions for a range of species at different trophic levels against the AA proportions found in the *Drosophila* exome-matched profile (Figure 5A). This included yeast (*Saccharomyces cerevisiae*), the fly’s food source (Markow, 2015), plants on which yeast may grow (*Zea mays, Solanum tuberosum*) as well as an organism that eats *Drosophila* (the spider *Parasteatoda tepidariorum*), and two additional levels of higher consumers (chicken *Gallus gallus* and humans *Homo sapiens*). We found that the AA profiles of the higher eukaryotes were more similar to the FLYAA profile than that of the autotroph yeast, as anticipated (Figure 5A). However, high similarity to FLYAA was not simply limited to AA heterotrophs, since the plants, which are AA autotrophs, were more similar to FLYAA than the similarity found between yeast and FLYAA. In summary, the AA proportions encoded by the exomes of heterotrophs in a trophic chain do indeed appear to be highly similar to each other as we predicted, but the usage of AAs amongst autotrophs can vary more widely.

**Figure 5.**
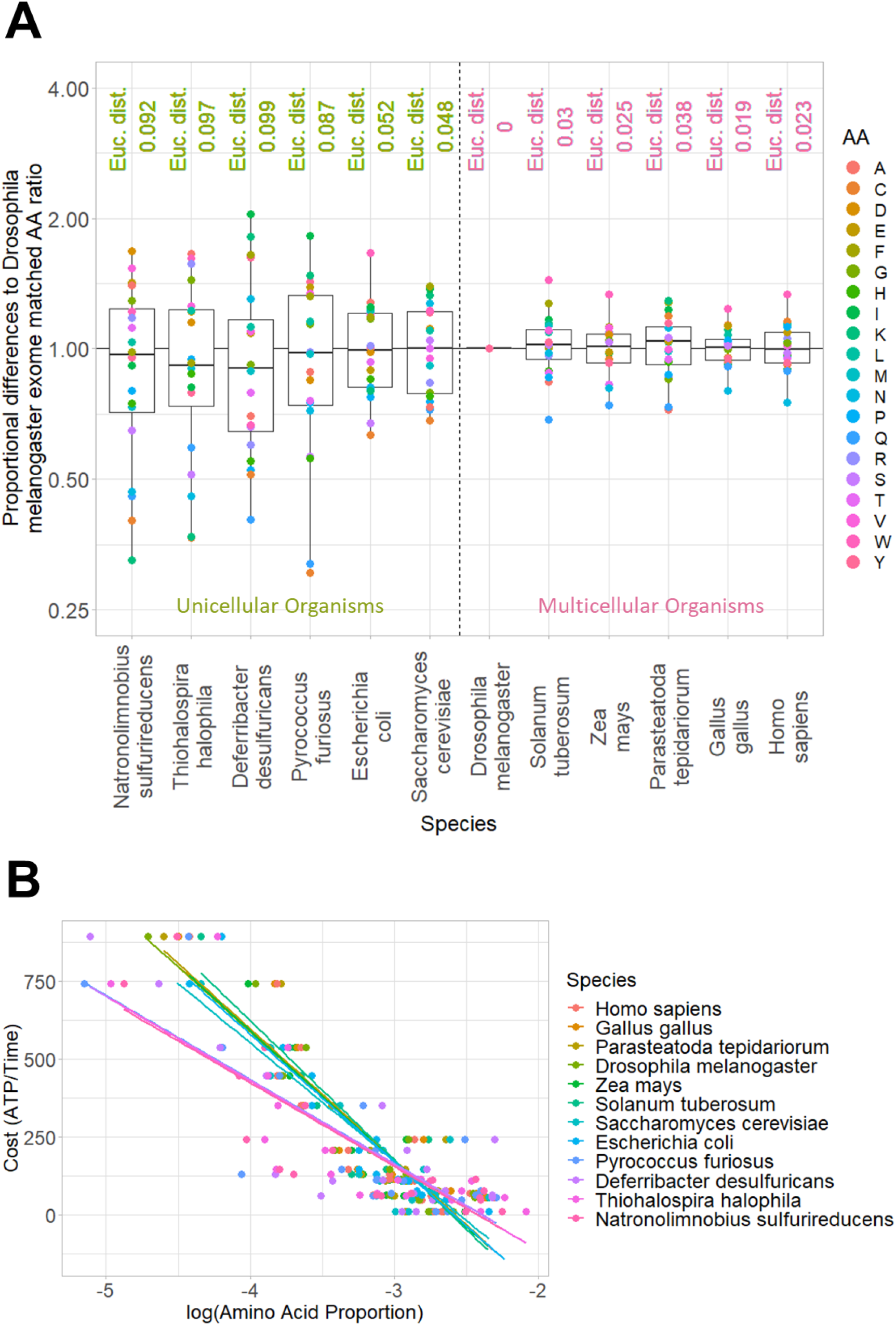
Cross-species comparisons of genome-wide AA proportions indicate constraints at the level of the energetic costs of biosynthesis. A) Proportional difference in AA concentrations between the exome matched AA ratio (FLYAA) and several multicellular (right) and unicellular (left) species. The Euclidean distance (indicated at the top of the plot) indicates the degree of divergence of each ratio from the exome of *Drosophila melanogaster*. B) The log-transformed exome matched AA proportions of all tested species and diets inversely correlated in a linear fashion with the metabolic cost of AA generation (R^2^≥0.67, which increased to R^2^≥0.79 in non-extremophiles). The equations derived from the linear models clustered in two groups based on their slope: extremophiles (*P. furiousus, D. desulfuricans, T. halophila, N. sulfurireducens*) (m = -270 ±1) and all other organisms (m = -416 ±18).

In an attempt to explain the pattern of AAs found for all species in our comparisons, we assessed the AA profiles as a function of their metabolic costs of biosynthesis, as described in (Krick *et al*., 2014). We found that conservation of the energetic costs of biosynthesis for each AA provides a good explanation for the relative abundance of each AA in the predicted proteome of each of the seven organisms in our trophic chain (autotrophs and heterotrophs) and that the slopes of this relationship between the organisms were indistinguishable (m = - 416 ±18, R^2^≥0.79) (Figure 5B). Thus, although the AA profile of yeast appeared to be more distant from the fly than were the plants, this divergence involved changes in AA usage that caused no net change in the predicted metabolic cost of producing the proteome. Thus, the relative proportion of AAs encoded by an organism’s exome can be explained by their metabolic costs of biosynthesis at the level of autotrophs, and this may constrain their pattern of usage in heterotrophs at higher trophic levels because they are limited by AA availability. This pattern of divergent AA profiles with preserved cost was also true when we extended the analysis to include another common unicellular AA autotroph, *Escherichia coli* (Figure 5B). However, when we also included several extremophiles in our analysis, two microbes that live in hydrothermal vents (*Deferribacter desulfuricans, Pyrococcus furiosus*) and two from salt lakes (*Natronolimnobius sulfurireducens, Thiohalospira halophila*), they continued to show a strong inverse relationship between the cost of biosynthesis and the relative abundance of each AA, but with a shallower slope (m = -270 ±1, R^2^≥0.67)(Figure 5B). Thus, environmental conditions may constrain the proteome profiles of producers according to the metabolic costs of AA production, and this may shape the pattern of AA usage across trophic levels from unicellular autotrophs to higher-order predators.

## DISCUSSION

Dietary nitrogen availability limits reproductive fitness and population growth in multiple species, from algae to insects, birds, and mammals (Smith, 1968, 1991; Nixon, McClain and Donohoe, 1975; Bosman and Hockey, 1986; White, 1993; Ågren, 2004; Bi *et al*., 2005; Webb *et al*., 2005; Grandison, Piper and Partridge, 2009). For higher organisms, this limitation has been shown to occur at the level of the single essential AA that is most undersupplied in the diet relative to physiological demands (Grandison, Piper and Partridge, 2009; Piper *et al*., 2017; Solon-Biet *et al*., 2019). To overcome this limitation, organisms have evolved numerous behavioural, symbiotic, biochemical, and physical adaptations that enable them to seek out, acquire, and metabolise dietary nitrogen and/or protein with high efficiency (White, 1993; Sørensen *et al*., 2008; Gosby *et al*., 2011; Simpson and Raubenheimer, 2012; Simpson, Le Couteur and Raubenheimer, 2015; Almeida de Carvalho and Mirth, 2017; Leitão-Gonçalves *et al*., 2017; Simpson *et al*., 2017; Solon-Biet *et al*., 2019). Our data here reveal that these systems achieve additional levels of efficiency because organisms require a relatively constant proportion of AAs across diverse conditions, and this proportion is closely matched to the diets they consume.

We observed that female and male flies show a marked difference in the absolute levels of dietary protein that is required to support maximal fecundity, with females requiring ∼5-times more than males. This difference is in agreement with females having a higher anabolic demand for producing gametes and is reflected in the fact that females express a greater preference for protein-rich diets than males (Camus *et al*., 2018). However, we were surprised to find that although males and females differ in expression of ∼50% of their genome, they exhibit no observable difference in proportions of AAs (i.e. protein quality) that they require to maximise fecundity. What’s more, the calculated transcriptome-weighted AA profiles of both males and females are similar to the AA profile calculated from the exome without including expression weightings. These data indicate that for other important physiological changes that require less substantial remodelling of the transcriptome, such as when launching an immune response, the flies’ requirements for each AA will be met by the same, exome-matched protein quality. For this reason, we predict that exome matching (without weighting for gene expression) is a generally good estimation of the flies’ dietary AA requirements under a broad range of conditions.

Despite the general conservation that we found in the exome-matched AA profiles for consumers at higher trophic levels, we observed an increased variability amongst AA autotrophs. This observation highlights a new nutritional dimension (to the level of AAs) to the already recognised divergence in the flexibility of biomass composition in autotrophs when compared to heterotrophic consumers (Sperfeld *et al*., 2017). It also means that animals consuming at the autotroph/heterotroph interface, such as flies feeding on their natural food source, yeast, are more likely to experience a mismatch between the profile of AAs they consume and what they require for fitness. As a consequence of this mismatch, we predict that organisms feeding at this interface are more likely to exhibit enhanced performance when their natural diets are supplemented with essential AAs than are heterotrophs that consume other heterotrophs (e.g. hyper carnivores). For several organisms, including flies, there is empirical evidence to support this prediction (Peoples *et al*., 1994; Ramsay and Houston, 1998; Webb *et al*., 2005; Grandison, Piper and Partridge, 2009; Piper *et al*., 2014). Further, if this mismatch is an ongoing fitness constraint, consumers should evolve mechanisms to specifically buffer against shortfalls in the supply of the most limiting AA (Anderson, Boersma and Raubenheimer, 2004). In other work, we have found that flies feeding on their natural food source, yeast, are methionine limited for egg production (Grandison, Piper and Partridge, 2009; Piper *et al*., 2017) and when we omit any one of the flies’ 10 essential AAs from the diet, egg-laying is arrested (Sang and King, 1961) (André Nogueira Alves, *personal communication*). Interestingly, the rate at which egg-laying declines varies depending on the identity of the missing AA: for the omission of any one of eight of the flies’ ten essential AAs, there is a rapid arrest in egg production that takes ∼3 days, whereas omitting either methionine or histidine results in a much slower decline, taking ∼7 days and ∼5 days respectively to cease egg production (André Nogueira Alves, *personal communication*). Thus, flies have evolved a mechanism that buffers egg production against fluctuations in the most limiting essential AA (methionine) as well as the AA that exome matching predicts to be the second most limiting (histidine). Interestingly, in other work, we have found that egg-laying of flies feeding on yeast is not only limited by dietary AAs, but also by dietary sterols (Zanco *et al*., 2021) - a nutritional co-limitation that has been recognised in other organisms (Martin-Creuzburg, Sperfeld and Wacker, 2009; Wacker and Martin-Creuzburg, 2012). Again, we find that flies can buffer against fluctuations in this limiting nutrient, since they continue to produce high-quality eggs even when the diet is completely depleted of sterols (Zanco *et al*., 2021). By contrast, flies quickly arrest egg production in the absence of any other essential micronutrient (Wu *et al*., 2020). The agreement between the identity of the nutrients against which the flies possess buffering capacity and those that we have found experimentally to be limiting in the diet makes a compelling case that these are specifically evolved adaptations.

Understanding the way organisms interact with their nutritional environments has been the topic of an enormous body of research. Historically, the models employed to study ecosystem dynamics have employed a single currency (energy), but work over the last 30 years has described and modelled the role of other nutritional components in ecological constraints and organismal fitness (Lindeman, 1991; Sterner and Elser, 2002; Simpson and Raubenheimer, 2012). Ecological Stoichiometry and Nutritional Geometry are two such models that provide complementary approaches to describing these systems (Sperfeld *et al*., 2017; Anderson *et al*., 2020; Burian, Nielsen and Winder, 2020). Ecological Stoichiometry explains individual and ecosystem-level phenomena as shaped by the availability of energy and the chemical elements (e.g. carbon, nitrogen and phosphorous) that make up biomass (Sterner and Elser, 2002). By contrast, Nutritional Geometry models the same phenomena from the point of view of the biochemicals (e.g. carbohydrates and proteins) that modify feeding behaviour and evolutionary fitness (Simpson and Raubenheimer, 1993). Our data show an association between the energetic and elemental constraints on autotrophs in shaping the proportions of the biochemicals (AAs) whose limitation shapes the composition of heterotroph biomass. These data underscore the importance of considering the most relevant constraints of the trophic level under study and highlight the need for emerging models that seek to combine the two approaches (Anderson, Boersma and Raubenheimer, 2004; Anderson *et al*., 2020; Burian, Nielsen and Winder, 2020).

Part of the motivation of our study was to generate a new way of tailoring dietary AA profiles to meet the changing demands of organisms as they progress through development and experience altered health. Here we find no theoretical or experimental evidence to undertake such measures since the organism’s AA requirements for its expressed genome tend to converge on that encoded by the exome. Thus, exome matching can serve as a simple guide to establish the dietary AA requirements of animals and so spare some of the expensive, time-consuming, and physically invasive efforts that are employed to determine the optimal dietary AA requirements of animals in agriculture (Levesque *et al*., 2010; Boye, Wijesinha-Bettoni and Burlingame, 2012) and for humans for health (Institute of Medicine, 2005).

## MATERIALS AND METHODS

### DIETS

#### Sugar/yeast (SY) food

Our sugar-yeast (SY) food contained sugar (Bundaberg, M180919) (50 g/L), brewer’s yeast (MP Biomedicals, 903312) (100 g/L), agar (Gelita, A-181017) (10 g/L), propionic acid (Merck, 8.00605.0500) (3 mL/L) and nipagin (Sigma-Aldrich, W271004-5KG-K) (12 g/L), prepared as in (Bass et al., 2007). Refer to Supplementary Table 1 for more detailed product information.

#### HOLIDIC DIETS

##### EXOME-MATCHED AA RATIO DIET (FLYAA)

Chemically defined (holidic) diets used the recipe and were prepared as in (2). In all cases, the concentration of all nutrients, except AAs, were held constant at the levels published in (Piper *et al*., 2017) (Supplementary Table 1 and 2). The AA ratio was either matched to the fly exome, FLYAA (Piper *et al*., 2017), or the transcriptome-weighted dietary AA ratios (Table 1). Refer to Supplementary Table 1 for more detailed product information.

**Table 1.** Molar ratio of AAs. The table shows the molar ratio of AAs in the exome matching diet (FLYAA) and in the male (MALEAA) and female (FEMALEAA) transcriptome matching diets.

|  |  | MOLAR RATIO |  |  |
| --- | --- | --- | --- | --- |
|  |  | MALEAA | FEMALEAA | FLYAA |
| ESSENTIAL AMINO ACIDS | **F** | 0.039 | 0.036 | 0.037 |
|  | **H** | 0.023 | 0.024 | 0.026 |
|  | **I** | 0.056 | 0.054 | 0.052 |
|  | **K** | 0.062 | 0.068 | 0.057 |
|  | **L** | 0.093 | 0.088 | 0.094 |
|  | **M** | 0.026 | 0.024 | 0.025 |
|  | **R** | 0.051 | 0.055 | 0.057 |
|  | **T** | 0.054 | 0.055 | 0.056 |
|  | **V** | 0.063 | 0.064 | 0.062 |
|  | **W** | 0.011 | 0.011 | 0.010 |
| NON-ESSENTIAL AMINO ACIDS | **A** | 0.079 | 0.078 | 0.075 |
|  | **C** | 0.019 | 0.016 | 0.017 |
|  | **D** | 0.049 | 0.051 | 0.053 |
|  | **E** | 0.059 | 0.062 | 0.063 |
|  | **G** | 0.068 | 0.068 | 0.062 |
|  | **N** | 0.047 | 0.047 | 0.047 |
|  | **P** | 0.052 | 0.049 | 0.052 |
|  | **Q** | 0.044 | 0.046 | 0.046 |
|  | **S** | 0.075 | 0.074 | 0.079 |
|  | **Y** | 0.030 | 0.030 | 0.031 |

### FLY STRAIN AND CONDITIONS

All experiments were carried out using our outbred wild-type strain of *Drosophila melanogaster* called Dahomey (Mair, Piper and Partridge, 2005). Stock and experimental flies were kept under controlled conditions of 25°C with at least 65% relative humidity for a 12:12 h photoperiod. Stocks were maintained on SY food.

### MALE FECUNDITY ASSAY

All flies were reared under standard density conditions as detailed in Bass et al., 2007 (Bass *et al*., 2007). Males and virgin female flies were collected from 0 to 5 h after emerging and kept separately in fresh standard SY food vials. When flies were two days old, male flies were transferred to chemically defined diets with different AA ratios (FLYAA or MALEAA) and different total AA concentrations (0 g/L, 1.1 g/L, 2.1 g/L or 10.7 g/L). Following a seven-day adaptation period, individually housed males were placed with ten new age-matched (+/- 1 day) virgin females. After being co-housed for 24h, the females were removed and replaced with another ten virgin females. This procedure was repeated every day for seven days. All females that were removed from the vial containing the male were singly housed in a new vial containing SY food. After ten days on SY food, the vials with singly housed females were checked for the presence of offspring. For each male, the number of females he fertilised during each of these seven days was recorded.

### FEMALE EGG-LAYING ASSAY

Upon emerging as adults, male and female flies were transferred to fresh SY food and kept together for 48hours. Then female flies were then separated from males and kept on chemically defined diets with different AA ratios (FLYAA or FEMALEAA) and different total AA concentrations (0 g/L, 5.4 g/L or 10.7 g/L). Each vial contained five females, and flies were transferred to fresh food every 24h for eight days. The numbers of eggs laid per vial per day for eight days were counted, using QuantiFly software (2.0), and recorded (Waithe *et al*., 2015).

### CALCULATING THE EXOME AND TRANSCRIPTOME MATCHED AA PROPORTIONS

The fly exome matched AA ratio (FLYAA) was calculated as in (Piper *et al*., 2017). Modifications to this procedure were performed as described below.

#### DEVELOPMENT OF DIETARY TRANSCRIPTOME MATCHED AA RATIOS

To estimate the transcriptome matched AA ratio, we computed the relative proportion (*P*) of each AA in the transcriptome, *P*(*AA*_*i*_) (where *i* indicates one of the 20 protein-coding AAs). This calculation was performed in two steps. First, we calculated the number of instances of each AA encoded by each protein isoform, *AA*_*ij*_ (where *j* indicates a protein isoform). This overcame the previous limitation in exome matching in which protein isoforms and length was not considered in the exome matched calculation.

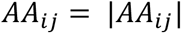

To generate the transcriptome-weighted values for each AA (*AA*_*i*_), we multiplied the number of each AA in each isoform, *AA*_*ij*_, by the isoform’s expression level, *E*_*j*_ (measured in FPKM levels). We then summed the transcriptome-weighted AA abundance for each AA for all protein isoforms in the expressed genome.

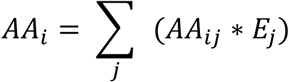

To obtain the final proportion of each AA encoded by the transcriptome weighted exome *P*(*AA*_*i*_), we divided the total number of each type of AA, *AA*_*i*_, by the sum of all transcriptome-weighted AAs, ∑_*i*_ *AA*_*i*_.

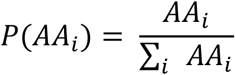

Thus, the sum of the proportions of all transcriptome-weighted AAs was equal to one.

The dietary transcriptome matched AA ratios for male and female flies were obtained as the average AA ratio of five male and five female fly transcriptomes, three belonging to FlyAtlas 2 and two to ModEncode database (Table 1).

For all bioinformatics processing, we used R (3.4.4) and the R packages “seqinr” (3.4-5) and “stringr” (1.3.1) (Charif and Lobry, 2007; Wickham, 2010).

#### IDENTIFICATION OF LIMITING AAS

The identification of the limiting AA, and the degree to which it was predicted limiting, was performed as described in (Piper *et al*., 2017).

#### TISSUE-SPECIFIC DATA COMPARISON

The AA ratio Cleveland plots generated to compare dietary AA ratios with tissue-specific transcriptome matched AA ratios were produced using the R package “ggplot2” (3.1.1) (Valero-Mora, 2010). On these Cleveland plots, we identified the limiting AA in the exome (FLYAA) and the whole male (MALEAA) and female (FEMALEAA) transcriptome matched dietary AA ratios assuming that the tissue AA demand corresponded to the tissue-specific transcriptome matched AA ratios. The coverage value of each AA was calculated by dividing the molar proportion of the AA supplied in each diet (FEMALEAA, MALEAA, or FLYAA) by the molar proportion of each tissue-specific AA ratio. This ratio was expressed as a percentage value. For every tissue, the essential AA with the lowest percentage coverage was predicted to be limiting.

#### AA BIOSYNTHETIC COSTS

AA generation costs were obtained from (Krick *et al*., 2014). Linear regression analysis between AA generation costs and the exome matching AA proportions from multiple species was performed using the “lm” (3.4.4) function in R and plotted using “ggplot2” (3.1.1) (Valero-Mora, 2010).

The exome matched AA ratio from various species was calculated by taking the sum of each AA’s usage when all genes in the exome are counted once (i.e. not transcriptome weighted).

#### TRANSCRIPTOME DATA PERMUTATIONS

To calculate the transcriptome matched AA ratio, we sum the number of each AA in each gene multiplied by its expression calculated by FPKM. Then we divide the total count for each AA across all genes by the total number of all AAs across all genes. For each of the 20,000 permutations, we performed the same calculation after randomly assigning the expression weightings to genes in the genome.

### TRANSCRIPTOMES

#### Files and databases

Transcriptomic data for wild-type *Drosophila melanogaster* were obtained from Flyatlas 2, (European Nucleotide Archive, ENA, Study Accession: PRJEB22205) and ModEncode (ENA Study Accession: SRP006203) (Robinson *et al*., 2013; Brown and Celniker, 2015). The transcriptomes downloaded corresponded to RNAseq files from whole male flies (Flyatlas 2, ModEnconde), whole female flies (Flyatlas 2, ModEncode), and tissue-specific RNAseq data from male and female flies (Flyatlas 2). Reference genomes, gene annotations, and translated exomes were downloaded from the Ensembl database and Flybase (release: r6.21).

#### Transcriptome processing

RNAseq raw reads were trimmed for low-quality bases by Trimmomatic (0.38) (Bolger, Lohse and Usadel, 2014) using default quality cutoff parameters (http://www.usadellab.org/cms/?page=trimmomatic). Any remaining rRNA reads were removed using SortmeRNA (4.2.0) (Kopylova, Noé and Touzet, 2012). Quality controls of the raw, trimmed, and aligned reads were performed by FastQC (0.11.17) and multiQC (v1.6) (Andrews *et al*., 2015; Ewels *et al*., 2016). After trimming rRNA and low-quality reads, at least 4 million reads were obtained on each trimmed RNAseq sample with at least a 90% overall alignment rate and a 75% unique alignment rate with the reference genome. Quality control data is available on request. Then, we used Hisat 2 (2.1.0) and Samtools (1.9), to map, index and sort the trimmed reads to the reference genome (Flybase release: r6.21) (Kim, Langmead and Salzberg, 2015). Transcript quantification was performed as Fragments per kilobase per million mapped read (FPKM) using Cufflinks (2.2.1) (Trapnell *et al*., 2010).

## DATA AVAILABILITY

The gene expression file was generated from RNASeq samples downloaded from the FlyAtlas 2 and modENCODE databases (see Materials and Methods).

## AUTHOR CONTRIBUTIONS

All authors contributed to the planning, performing, and/or analysis of research. All authors suggested experiments, reviewed, and edited the manuscript. J.G.O. and M.D.W.P drafted the manuscript.

## ACKNOWLEDGEMENTS

We thank Amy Dedman for her aid in research and laboratory management. We acknowledge the funding of the ARC (FT150100237) and the NHMRC (1182330) to M.D.W.P., as well as the ARC (FT170100259) to C.K.M.

## SUPPLEMENTARY MATERIAL

Supplementary Table 1. Detailed information of dietary components.

Supplementary Table 2. Composition of chemically defined diets.

## Notes

### Competing Interest Statement

The authors have declared no competing interest.

## REFERENCES

Ågren, G. I. (2004) ‘The C:N:P stoichiometry of autotrophs - Theory and observations’, Ecology Letters, 7(3), pp. 185–191. doi: 10.1111/j.1461-0248.2004.00567.x.

Almeida de Carvalho, M.J. and Mirth, C. K. (2017) ‘Food intake and food choice are altered by the developmental transition at critical weight in Drosophila melanogaster’, Animal Behaviour. Academic Press, 126, pp. 195–208. doi: 10.1016/j.anbehav.2017.02.005.

Anderson, T. R. et al. (2020) ‘Geometric Stoichiometry: Unifying Concepts of Animal Nutrition to Understand How Protein-Rich Diets Can Be “Too Much of a Good Thing”‘, Frontiers in Ecology and Evolution. Frontiers Media S.A., 8, p. 196. doi: 10.3389/fevo.2020.00196.

Anderson, T. R., Boersma, M. and Raubenheimer, D. (2004) ‘Stoichiometry: Linking elements to biochemicals’, Ecology. Ecological Society of America, 85(5), pp. 1193–1202. doi: 10.1890/02-0252.

Andrews, S. et al. (2015) ‘FastQC. A quality control tool for high throughput sequence data. Babraham Bioinformatics’, Babraham Institute, 1(1), p. 1. Available at: http://www.bioinformatics.babraham.ac.uk/projects/fastqc/.

Bass, T.M. et al. (2007) ‘Optimization of Dietary Restriction Protocols in Drosophila’, The Journals of Gerontology: Series A, 62(10), pp. 1071–1081. doi: 10.1093/gerona/62.10.1071.

Bi, J. L. et al. (2005) ‘Influence of seasonal nitrogen nutrition fluctuations in orange and lemon trees on population dynamics of the glassy-winged sharpshooter (Homalodisca coagulata)’, Journal of Chemical Ecology. Springer, 31(10), pp. 2289–2308. doi: 10.1007/s10886-005-7102-3.

Bolger, A. M., Lohse, M. and Usadel, B. (2014) ‘Trimmomatic: A flexible trimmer for Illumina sequence data’, Bioinformatics. Oxford University Press, 30(15), pp. 2114–2120. doi: 10.1093/bioinformatics/btu170.

Bosman, A. and Hockey, P. (1986) ‘Seabird guano as a determinant of rocky intertidal community structure’, Marine Ecology Progress Series, 32, pp. 247–257. doi: 10.3354/meps032247.

Boye, J., Wijesinha-Bettoni, R. and Burlingame, B. (2012) ‘Protein quality evaluation twenty years after the introduction of the protein digestibility corrected AA score method’, British Journal of Nutrition. Br J Nutr, 108(SUPPL. 2). doi: 10.1017/S0007114512002309.

Brown, J. B. and Celniker, S. E. (2015) ‘Lessons from modENCODE’, Annual Review of Genomics and Human Genetics, 16(1), pp. 31–53. doi: 10.1146/annurev-genom-090413-025448.

Burian, A., Nielsen, J. M. and Winder, M. (2020) ‘Food quantity–quality interactions and their impact on consumer behavior and trophic transfer’, Ecological Monographs. Ecological Society of America, 90(1), p. e01395. doi: 10.1002/ecm.1395.

Camus, M. F. et al. (2018) ‘Dietary choices are influenced by genotype, mating status, and sex in Drosophila melanogaster’, Ecology and Evolution. John Wiley and Sons Ltd, 8(11), pp. 5385–5393. doi: 10.1002/ece3.4055.

Camus, M. F., Piper, M. D. W. and Reuter, M. (2019) ‘Sex-specific transcriptomic responses to changes in the nutritional environment’, eLife. eLife Sciences Publications Ltd, 8. doi: 10.7554/eLife.47262.

Charif, D. and Lobry, J. R. (2007) ‘SeqinR 1.0-2: A Contributed Package to the R Project for Statistical Computing Devoted to Biological Sequences Retrieval and Analysis’, in, pp. 207–232. doi: 10.1007/978-3-540-35306-5_10.

Denno, R. F. and Fagan, W. F. (2003) ‘Might nitrogen limitation promote omnivory among carnivorous arthropods?’, Ecology. Ecological Society of America, pp. 2522–2531. doi: 10.1890/02-0370.

Ewels, P. et al. (2016) ‘MultiQC: summarize analysis results for multiple tools and samples in a single report’, Bioinformatics, 32(19), pp. 3047–3048. doi: 10.1093/bioinformatics/btw354.

Glück, E. (1988) ‘Why do parent birds swallow the feces of their nestlings?’, Experientia. Birkhäuser-Verlag, 44(6), pp. 537–539. doi: 10.1007/BF01958943.

Gosby, A.K. et al. (2011) ‘Testing protein leverage in lean humans: A randomised controlled experimental study’, PLoS ONE. PLoS One, 6(10). doi: 10.1371/journal.pone.0025929.

Grandison, R. C., Piper, M. D. W. and Partridge, L. (2009) ‘Amino-acid imbalance explains extension of lifespan by dietary restriction in Drosophila’, Nature. Nature Publishing Group, 462(7276), pp. 1061–1064. doi: 10.1038/nature08619.

Hegedus, Z. et al. (2009) ‘Deep sequencing of the zebrafish transcriptome response to mycobacterium infection’, Molecular Immunology. Pergamon, 46(15), pp. 2918–2930. doi: 10.1016/j.molimm.2009.07.002.

Institute of Medicine (2005) ‘Dietary Reference Intakes for Energy, Carbohydrate, Fiber, Fat, Fatty Acids, Cholesterol, Protein, and AAs (Macronutrients)’, National Academies Press, pp. 1–1331. doi: 10.17226/10490.

Jang, T. and Lee, K. P. (2018) ‘Comparing the impacts of macronutrients on life-history traits in larval and adult Drosophila melanogaster: The use of nutritional geometry and chemically defined diets’, Journal of Experimental Biology. Company of Biologists Ltd, 221(21). doi: 10.1242/jeb.181115.

Keith, M. O. and Bell, J. M. (1989) ‘The Utilization of Nitrogen for Growth in Mice Fed Blends of Purified Proteins’, Proceedings of the Society for Experimental Biology and Medicine. Proc Soc Exp Biol Med, 190(3), pp. 246–253. doi: 10.3181/00379727-190-42856.

Kim, D., Langmead, B. and Salzberg, S. L. (2015) ‘HISAT: A fast spliced aligner with low memory requirements’, Nature Methods. Nature Publishing Group, 12(4), pp. 357–360. doi: 10.1038/nmeth.3317.

Kopylova, E., Noé, L. and Touzet, H. (2012) ‘SortMeRNA: Fast and accurate filtering of ribosomal RNAs in metatranscriptomic data’, Bioinformatics, 28(24), pp. 3211–3217. doi: 10.1093/bioinformatics/bts611.

Krick, T. et al. (2014) ‘AA metabolism conflicts with protein diversity’, Molecular Biology and Evolution. Oxford University Press, 31(11), pp. 2905–2912. doi: 10.1093/molbev/msu228.

Leitão-Gonçalves, R. et al. (2017) ‘Commensal bacteria and essential AAs control food choice behavior and reproduction’, PLoS Biology. Public Library of Science, 15(4), p. e2000862. doi: 10.1371/journal.pbio.2000862.

Levesque, C. L. et al. (2010) ‘Review of advances in metabolic bioavailability of AAs’, Livestock Science, pp. 4–9. doi: 10.1016/j.livsci.2010.06.013.

Li, S. T. et al. (2019) ‘DAF-16 stabilizes the aging transcriptome and is activated in mid-aged Caenorhabditis elegans to cope with internal stress’, Aging Cell. Blackwell Publishing Ltd, 18(3), p. e12896. doi: 10.1111/acel.12896.

Lindeman, R. L. (1991) ‘The trophic-dynamic aspect of ecology’, Bulletin of Mathematical Biology. Kluwer Academic Publishers, 53(1–2), pp. 167–191. doi: 10.1007/BF02464428.

Lochmiller, R. L. et al. (1995) ‘Habitat-induced changes in essential amino-acid nutrition in populations of eastern cottontails’, Journal of Mammalogy. Allen Press Inc., 76(4), pp. 1164–1177. doi: 10.2307/1382608.

Mair, W., Piper, M. D. W. and Partridge, L. (2005) ‘Calories do not explain extension of life span by dietary restriction in Drosophila’, PLoS Biology. Public Library of Science, 3(7), pp. 1305–1311. doi: 10.1371/journal.pbio.0030223.

Manzoni, C. et al. (2018) ‘Genome, transcriptome and proteome: The rise of omics data and their integration in biomedical sciences’, Briefings in Bioinformatics, 19(2). doi: 10.1093/BIB/BBW114.

Markow, T. A. (2015) ‘The secret lives of Drosophila flies’, eLife. eLife Sciences Publications Ltd, 4(June). doi: 10.7554/eLife.06793.

Martin-Creuzburg, D., Sperfeld, E. and Wacker, A. (2009) ‘Colimitation of a freshwater herbivore by sterols and polyunsaturated fatty acids’, Proceedings of the Royal Society B: Biological Sciences. Royal Society, 276(1663), pp. 1805–1814. doi: 10.1098/rspb.2008.1540.

May, C. M. and Zwaan, B. J. (2017) ‘Relating past and present diet to phenotypic and transcriptomic variation in the fruit fly.’, BMC genomics. BioMed Central, 18(1), p. 640. doi: 10.1186/s12864-017-3968-z.

Moskalev, A. A. et al. (2019) ‘Transcriptome Analysis of Long-lived Drosophila melanogaster E(z) Mutants Sheds Light on the Molecular Mechanisms of Longevity’, Scientific Reports. Nature Publishing Group, 9(1). doi: 10.1038/s41598-019-45714-x.

Nixon, C. M. (1970) ‘Insects as Food for Juvenile Gray Squirrels’, American Midland Naturalist. JSTOR, 84(1), p. 283. doi: 10.2307/2423756.

Nixon, C. M., McClain, M. W. and Donohoe, R. W. (1975) ‘Effects of Hunting and Mast Crops on a Squirrel Population’, The Journal of Wildlife Management. JSTOR, 39(1), p. 1. doi: 10.2307/3800460.

Oshlack, A. and Wakefield, M. J. (2009) ‘Transcript length bias in RNA-seq data confounds systems biology’, Biology Direct 2009 4:1. BioMed Central, 4(1), pp. 1–10. doi: 10.1186/1745-6150-4-14.

Peoples, A. D. et al. (1994) ‘Limitations of AAs in diets of northern bobwhites (Colinus virginianus)’, American Midland Naturalist. University of Notre Dame, 132(1), pp. 104–116. doi: 10.2307/2426205.

Pible, O. and Armengaud, J. (2015) ‘Improving the quality of genome, protein sequence, and taxonomy databases: A prerequisite for microbiome meta-omics 2.0’, Proteomics, 15(20). doi: 10.1002/pmic.201500104.

Piper, M. D. W. et al. (2014) ‘A holidic medium for Drosophila melanogaster’, Nature Methods. Europe PMC Funders, 11(1), pp. 100–105. doi: 10.1038/nmeth.2731.

Piper, M. D. W. et al. (2017) ‘Matching Dietary AA Balance to the In Silico-Translated Exome Optimizes Growth and Reproduction without Cost to Lifespan’, Cell Metabolism. Elsevier, 25(3), pp. 610–621. doi: 10.1016/j.cmet.2017.02.005.

Ramsay, S. L. and Houston, D. C. (1998) ‘The effect of dietary AA composition on egg production in blue tits’, Proceedings of the Royal Society B: Biological Sciences, 265(1404), pp. 1401–1405. doi: 10.1098/rspb.1998.0448.

Robinson, S. W. et al. (2013) ‘FlyAtlas: database of gene expression in the tissues of Drosophila melanogaster.’, Nucleic acids research. Oxford University Press, 41(Database issue), pp. D744–50. doi: 10.1093/nar/gks1141.

Sang, J. H. and King, R. C. (1961) ‘Nutritional Requirements of Axenically Cultured Drosophila Melanogaster Adults’, Journal of Experimental Biology, 38(4), pp. 793–809. doi: 10.1242/jeb.38.4.793.

Simpson, S. J. et al. (2017) ‘Dietary protein, aging and nutritional geometry’, Ageing Research Reviews. Elsevier Ireland Ltd, pp. 78–86. doi: 10.1016/j.arr.2017.03.001.

Simpson, S. J., Le Couteur, D. G. and Raubenheimer, D. (2015) ‘Putting the balance back in diet’, Cell. Cell Press, pp. 18–23. doi: 10.1016/j.cell.2015.02.033.

Simpson, S. J. and Raubenheimer, D. (1993) ‘A multi-level analysis of feeding behaviour: The geometry of nutritional decisions’, Philosophical Transactions of the Royal Society B: Biological Sciences. Royal Society, 342(1302), pp. 381–402. doi: 10.1098/rstb.1993.0166.

Simpson, S. J. and Raubenheimer, D. (2012) ‘The nature of nutrition: A unifying framework from animal adaptation to human obesity’, Princeton University Press, pp. 1–239. doi: 10.5860/choice.50-2662.

Sjøberg, K. A. et al. (2020) ‘Effects of Short-Term Dietary Protein Restriction on Blood AA Levels in Young Men’, Nutrients. MDPI AG, 12(8), p. 2195. doi: 10.3390/nu12082195.

Smith, C. C. (1968) ‘The Adaptive Nature of Social Organization in the Genus of Three Squirrels Tamiasciurus’, Ecological Monographs. Wiley, 38(1), pp. 31–64. doi: 10.2307/1948536.

Smith, T. B. (1991) ‘Behavioural Ecology of the Galah Ecolophus Roseicapillus in the Wheatbelt of Western Australia. Ian Rowley’, The Quarterly Review of Biology, 66(4), pp. 504–504. doi: 10.1086/417390.

Solon-Biet, S. M. et al. (2015) ‘Macronutrient balance, reproductive function, and lifespan in aging mice’, Proceedings of the National Academy of Sciences of the United States of America. National Academy of Sciences, 112(11), pp. 3481–3486. doi: 10.1073/pnas.1422041112.

Solon-Biet, S. M. et al. (2019) ‘Branched-chain AAs impact health and lifespan indirectly via AA balance and appetite control’, Nature Metabolism. Nature Research, 1(5), pp. 532–545. doi: 10.1038/s42255-019-0059-2.

Sørensen, A. et al. (2008) ‘Protein-leverage in mice: The Geometry of macronutrient balancing and consequences for fat deposition’, Obesity. Obesity (Silver Spring), 16(3), pp. 566–571. doi: 10.1038/oby.2007.58.

Sperfeld, E. et al. (2017) ‘Bridging Ecological Stoichiometry and Nutritional Geometry with homeostasis concepts and integrative models of organism nutrition’, Functional Ecology. Edited by J. Harwood. Blackwell Publishing Ltd, 31(2), pp. 286–296. doi: 10.1111/1365-2435.12707.

Sterner, R. W. and Elser, J. J. (2002) ‘Ecological Stoichiometry: The Biology of Elements from Molecules to the Biosphere’, Princeton University Press, Princeton, New Jersey, USA. Princeton University Press, p. 439.

Trapnell, C. et al. (2010) ‘Transcript assembly and quantification by RNA-Seq reveals unannotated transcripts and isoform switching during cell differentiation’, Nature Biotechnology. Nature Publishing Group, 28(5), pp. 511–515. doi: 10.1038/nbt.1621.

Valero-Mora, P. M. (2010) ‘ggplot2: Elegant Graphics for Data Analysis’, Journal of Statistical Software. Springer International Publishing (Use R!), 35(Book Review 1), pp. 245–246. doi: 10.18637/jss.v035.b01.

Wacker, A. and Martin-Creuzburg, D. (2012) ‘Biochemical nutrient requirements of the rotifer Brachionus calyciflorus: Co-limitation by sterols and AAs’, Functional Ecology. John Wiley & Sons, Ltd, 26(5), pp. 1135–1143. doi: 10.1111/j.1365-2435.2012.02047.x.

Waithe, D. et al. (2015) ‘QuantiFly: Robust trainable software for automated Drosophila egg counting’, PLoS ONE. Public Library of Science, 10(5). doi: 10.1371/journal.pone.0127659.

Webb, R. E. et al. (2005) ‘Impact of food supplementation and methionine on high densities of cotton rats: Support of the amino-acid-quality hypothesis?’, Journal of Mammalogy. Oxford Academic, 86(1), pp. 46–55. doi: 10.1644/1545-1542(2005)086<0046:IOFSAM>2.0.CO;2.

White, T. C. R. (1993) ‘The Inadequate Environment’, Springer Berlin Heidelberg. Berlin, Heidelberg: Springer Berlin Heidelberg. doi: 10.1007/978-3-642-78299-2.

Wickham, H. (2010) ‘Stringr: Modern, consistent string processing’, R Journal, 2(2), pp. 38–40. doi: 10.32614/rj-2010-012.

Wilder, S. M. et al. (2013) ‘Arthropod food webs become increasingly lipid-limited at higher trophic levels’, Ecology Letters. Edited by F. Jordan. Blackwell Publishing Ltd, 16(7), pp. 895–902. doi: 10.1111/ele.12116.

Wu, Q. et al. (2020) ‘Sexual dimorphism in the nutritional requirement for adult lifespan in Drosophila melanogaster’, Aging Cell. Blackwell Publishing Ltd, 19(3), p. e13120. doi: 10.1111/acel.13120.

Zanco, B. et al. (2021) ‘A dietary sterol trade-off determines lifespan responses to dietary restriction in drosophila melanogaster females’, eLife. eLife Sciences Publications Ltd, 10, pp. 1–20. doi: 10.7554/eLife.62335.

